## Supplementary Table 1 and Figures 1-6 for "RAPSYN-Mediated Neddylation of BCR-ABL Alternatively Determines the Fate of Philadelphia Chromosome-positive Leukemia"

1. Supplementary Table S1
2. Supplementary Figures S1-S6

**Supplementary Table S1 Sequences of shRNA and primers**

| Name | Gene symbol (ID) | NCBI Reference Sequence | Sequence |
|---|---|---|---|
| shRNA targeting sequence: RAPSYN #1 | RAPSYN (5913) | NM_005055 .5 | GCATTGCAGGTGTGGACAAAG |
| shRNA targeting sequence: RAPSYN #2 | | | GGAGTGTTGTGAGGAGTCTAT |
| shRNA targeting sequence: RAPSYN #3 | | | TGCACGCCAGAGGCCCATTTA |
| shRNA targeting sequence: SRC #1 | SRC (6714) | NM_005417 .5 | GAGGGACCCTTCGAGATCATCACTT |
| shRNA targeting sequence: SRC #2 | | | CATCCTCAGGAACCAACAATT |
| shRNA targeting sequence: SRC #3 | | | CAGGTGTGGAGAGAGAGGCTTCAAT |
| shRNA targeting sequence: SRC #4 | | | GCGTCCATATTTAACATGTAA |
| shRNA targeting sequence: SRC #5 | | | GGTTGTAAATACTTTGCATATTGTC |
| h. RAPSYN RT-PCR Primers | RAPSYN (5913) | NM_005055 .5 | Forward: CGCTACAAGGAGATGCTGAAG |
| | | | Reverse: CTTGCAGTAGGAGATGGTCTTG |
| h. ACTIN RT-PCR Primers | ACTB (60) | NM_001101 .5 | Forward: CTACAATGAGCTGCGTGTGGC |
| | | | Reverse: CAGGTCCAGACGCAGGATGGC |
| h. GAPDH RT-PCR Primers | GAPDH (2597) | NM_002046 .7 | Forward: GCGTGACATTAAGGAGAAG |
| | | | Reverse: GAAGGAAGGCTGGAAGAG |

| | | | |
|---|---|---|---|
| HA-NEDD8 △GG mutant primers | NEDD8 (4738) | NM_006156.3 | Forward: GTTGGCTCTGAGAGCAGCTGGTGGTCTTAGGC |
| | | | Reverse: GCCTAAGACCACCAGCTGCTCTCAGAGCCAAC |
| GFP-RAPSYN C366A mutant primers | RAPSYN (5913) | NM_005055.5 | Forward: TCTACTGCGGCCTGGCCGGCGAGTCCATAG |
| | | | Reverse: CTATGGACTCGCCGGCCAGGCCGCAGTAGA |
| GST-RAPSYN Y59F mutant primers | | | Forward: GGAGATGGGCCGCTTTAAGGAGATGCTGAAGTT |
| | | | Reverse: AACTTCAGCATCTCCTTAAAGCGGCCCATCTCC |
| GST-RAPSYN Y152F mutant primers | | | Forward: GAGAAGGCCCTGCGCTTTGCCCACAACAATGAT |
| | | | Reverse: ATCATTGTTGTGGGCAAAGCGCAGGGCCTTCTC |
| GST-RAPSYN Y336F mutant primers | | | Forward: TGAGCGAGAGCATTTTTCGCAGCAAAGGGCTC |
| | | | Reverse: GCAGCCCTTTGCTGCGAAAAATGCTCTCGCTCA |
| His-BCR-ABL K257R mutant primers | | | Forward: CCCCCGCTTCCTGAGGGACAACCTGATCGAC |
| | | | Reverse: GTCGATCAGGTTGTCCCTCAGGAAGCGGGGG |
| His-BCR-ABL K500R mutant primers | | | Forward: GGCTTGGAGATGAGAAGATGGGTCCTGTCGGG |
| | | | Reverse: CCCGACAGGACCCATCTTCTCATCTCCAAGCC |

| | | | |
|---|---|---|---|
| His-BCR-ABL K739R mutant primers | | | Forward: CTGCTTCTCTGCACCAGGCTCAAGAAGCAGAGC |
| | | | Reverse: GCTCTGCTTCTTGAGCCTGGTGCAGAGAAGCG |
| His-BCR-ABL K802R mutant primers | | | Forward: GACATCCAGAGAGAGAGGAGGGCGAACAAGGC |
| | | | Reverse: GCCCTTGTTCGCCCTCCTCTCTCTCTGGATGTC |
| His-BCR-ABL K1025R mutant primers | | | Forward: GTCAACAGTCTGGAGAGACACTCCTGGTACCAT |
| | | | Reverse: ATGGTACCAGGAGTGTCTCTCCAGACTGTTGAC |
| His-BCR-ABL K1135R mutant primers | | | Forward: TCCCCCAACTACGACAGGTGGGAGATGGAACC |
| | | | Reverse: GCGTTCCATCTCCCACCTGTCGTAGTTGGGGGA |
| His-BCR-ABL K1590R mutant primers | | | Forward: CCCACCTGTGGAAGAGGTCCAGCACGCTGAC |
| | | | Reverse: GTCAGCGTGCTGGACCTCTTCCACAGGTGGG |
| His-BCR-ABL K1990R mutant primers | | | Forward: CGAGAGGCCATCAACAGACTGGAGAATAATCTC |
| | | | Reverse: GAGATTATTCTCCAGTCTGTTGATGGCCTCTCG |

Supplementary Figure 1

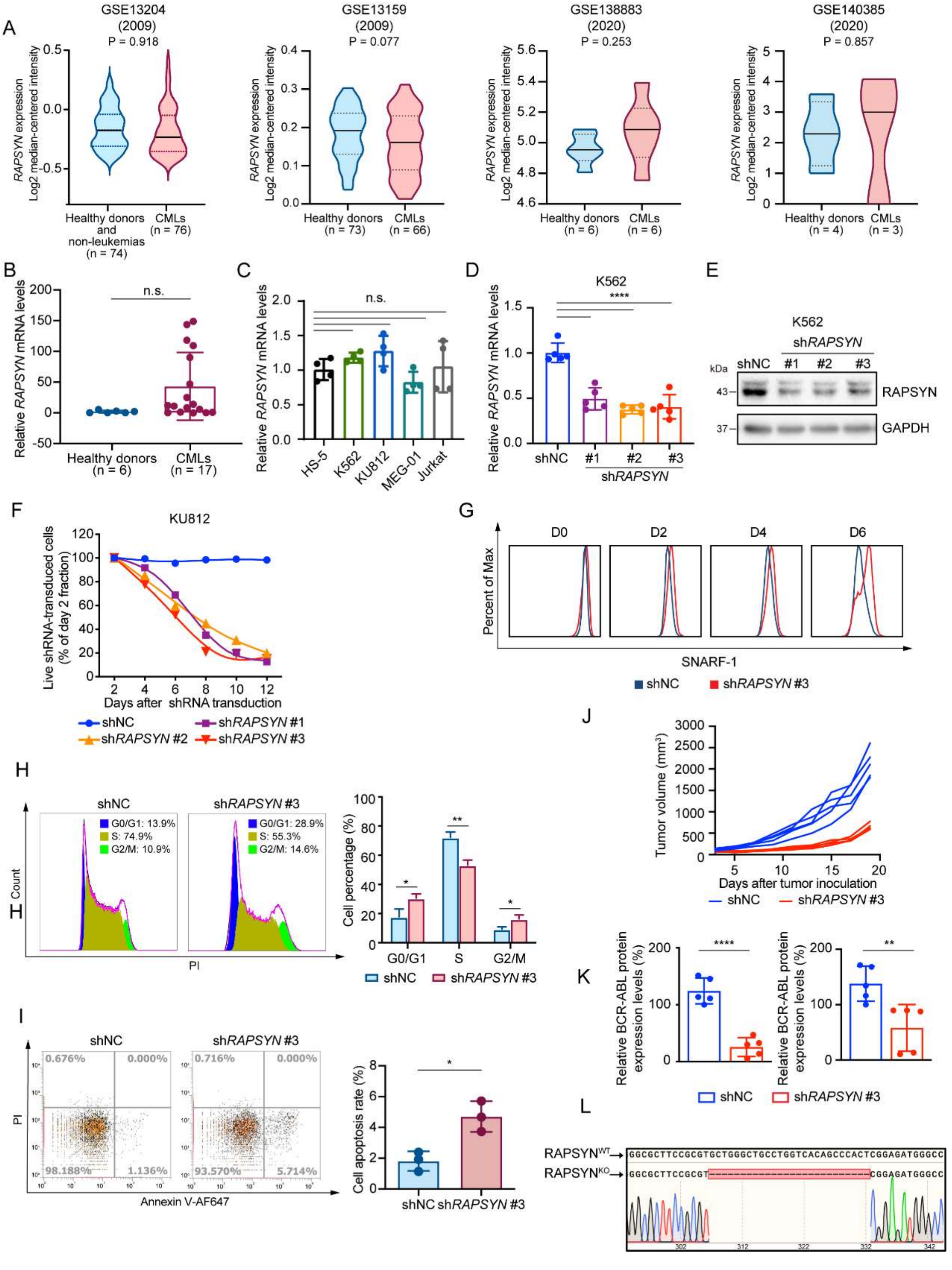


**Supplementary Figure 1. The mRNA levels of *RAPSYN* are not changed by Ph⁺ leukemia, whereas inhibition of RAPSYN suppresses Ph⁺ leukemia progression**

**A,** Analyses of *RAPSN* mRNA levels in PBMCs of healthy donors and non-leukemic patients compared to those of patients with CML from GSE13204, GSE13159, GSE138883 and GSE140385 datasets. **B,** Quantification of *RAPSYN* mRNA levels in

PBMCs of healthy donors and patients with $Ph^+$ leukemia from the cohort of primary samples using RT-PCR. **C,** Quantification of *RAPSYN* mRNA levels in $Ph^+$ leukemia cells (K562, KU812, MEG-01 and Jurkat) compared to normal bone marrow stromal cells (HS-5) using RT-PCR. **D,** Quantification of *RAPSYN* mRNA levels in K562 cells transduced with shNC or three independent shRNAs targeting *RAPSN* using RT-PCR. **E,** Immunoblotting of RAPSYN in K562 cells transduced with shNC or three different shRNAs targeting *RAPSN*. **F,** Cytotoxicity induced by shRNA-mediated *RAPSN* knockdown in KU812 cells. Representative results from at least 3 independent experiments are shown. **G,** Analysis of SNARF-1 labeling intensity in K562 cells transducted with shNC or sh*RAPSN* #3. **H,** Representative FACS cell cycle profiles of K562 cells transduced with shNC or sh*RAPSYN* #3. **I,** Representative FACS blots of apoptosis analysis of K562 cells transduced with shNC or sh*RAPSYN* #3. **J,** Individual growth curves of subcutaneous xenograft tumors were measured every two days from the third day after tumor inoculation for 19 days. **K,** Quantification of RAPSYN and BCR-ABL expression in mouse xenograft tumor biopsies from K562 cells transduced with sh*RAPSYN* #3 or shNC. **L,** Verification of $RAPSYN^{KO}$ in K562 cells. The red dotted line indicates deleted sequences. *RAPSN* mRNA levels were normalized to that of *ACTIN* (B) or *GAPDH* (C-D); error bars, mean ± SD; * $p < 0.05$, ** $p < 0.01$, **** $p < 0.0001$, n.s., not significant; unpaired Student's t-test (A-B, and J) or one-way ANOVA test (C-D).

Supplementary Figure 2

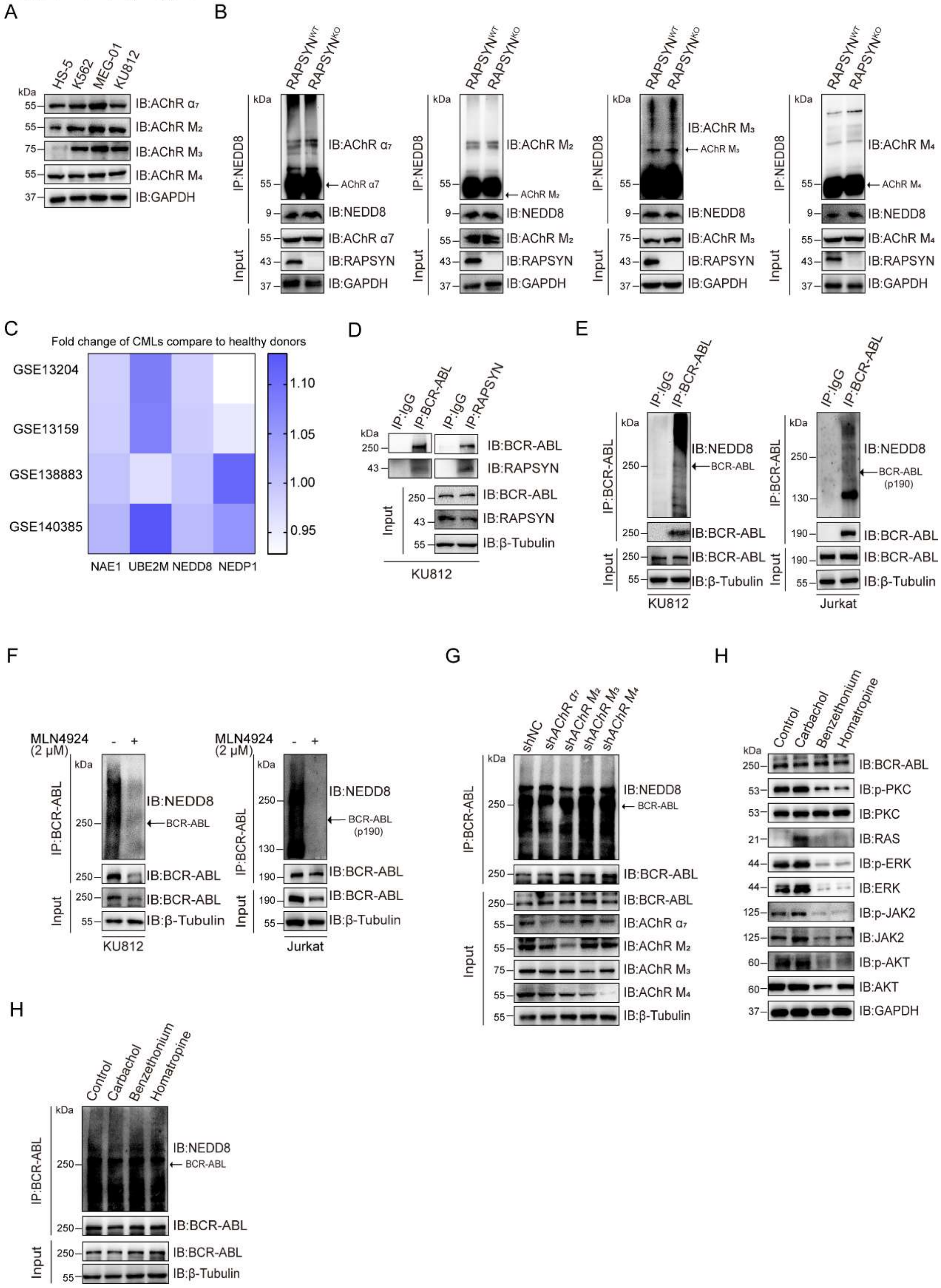


**Supplementary Figure S2. RAPSYN is an E3 ligase to neddylate BCR-ABL.**

**A,** Immunoblots of AChR subunits $\alpha_7$, $M_2$, $M_3$ and $M_4$ in Ph[+] leukemia cells (K562, KU812, and MEG-01) compared to normal bone marrow stromal cells (HS-5). **B,** Immunoblotting analyses of AChR subunit $\alpha_7$, $M_2$, $M_3$ and $M_4$ neddylation levels after immunoprecipitation of NEDD8 in WT and RAPSYN/KO K562 cells. **C,** Heatmap showing the fold change in mRNA level of neddylation-related proteins in CML patients compared to that in healthy donors. **D,** Immunoblots of BCR-ABL and RAPSYN in KU812 cells after co-immunoprecipitation with RAPSYN and BCR-ABL antibodies, respectively. **E,** Immunoblotting analyses of BCR-ABL neddylation after immunoprecipitation of BCR-ABL in KU812 and Jurkat cells. **F,** Immunoblotting analyses of BCR-ABL neddylation after immunoprecipitation of BCR-ABL in KU812 and Jurkat cells treated with MLN4924 or DMSO for 24 h. **G,** Immunoblotting analyses of BCR-ABL neddylation after immunoprecipitation of BCR-ABL in K562 cells transduced with sh$\alpha_7$, sh$M_2$, sh$M_3$, sh$M_4$ or shNC. **H,** Immunoblotting analyses of PKC-RAS-ERK and JAK2-AKT changes in K562 cells treated with AChR agonist carbamylcholine chloride (carbachol, 100 μM) and antagonist benzethonium (5 μM) for nAChR or tomatropine bromide (Homatropine, 5 μM) for mAChR for 24 h. **I,** Immunoblotting analyses of BCR-ABL neddylation after immunoprecipitation of BCR-ABL in K562 cells treated with AChR agonist carbamylcholine chloride (carbachol, 100 μM) and antagonist benzethonium (5 μM) for nAChR or tomatropine bromide (Homatropine, 5 μM) for mAChR for 24 h.

Supplementary Figure 3

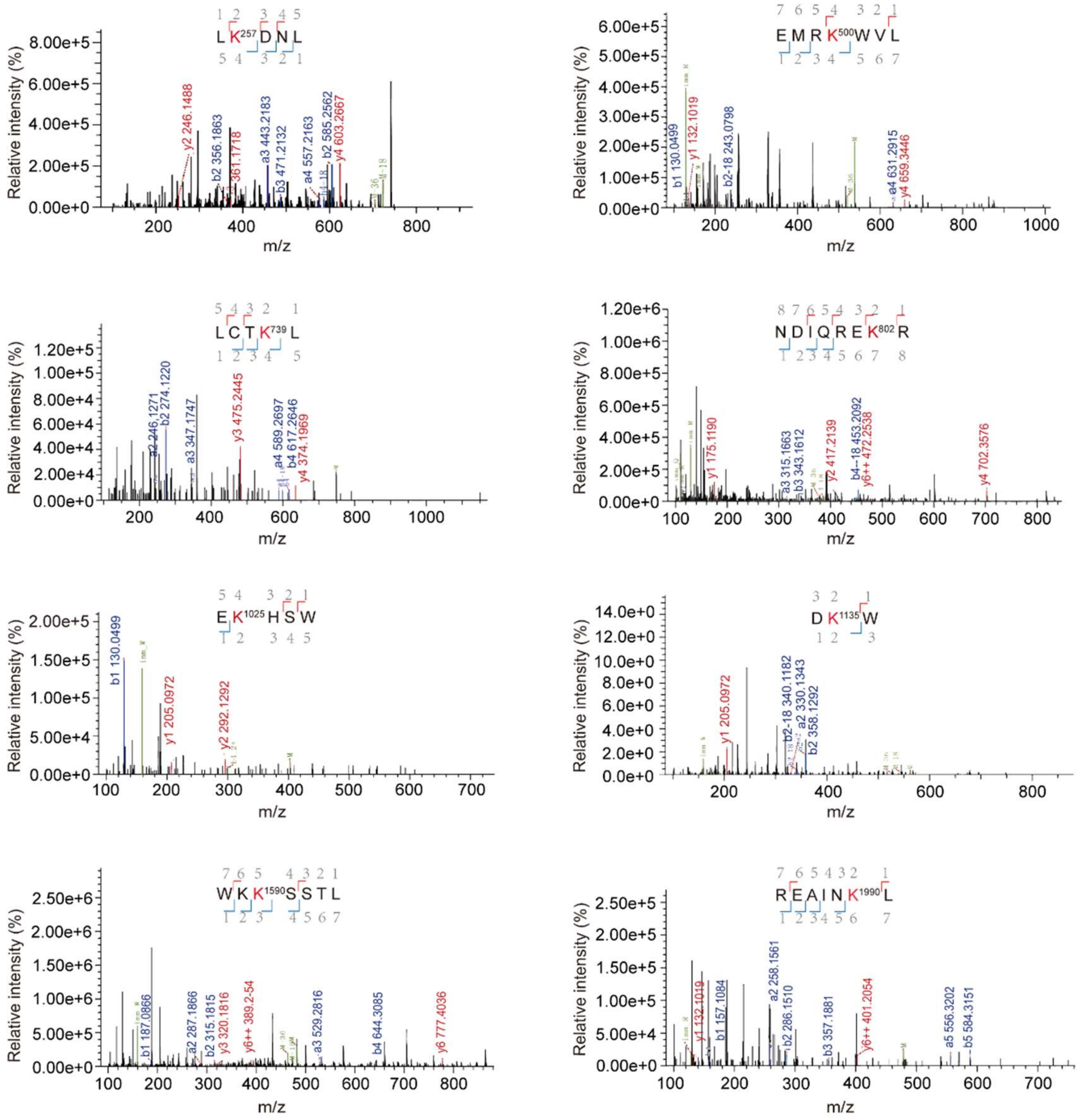


**Supplementary Figure S3. LC-MS/MS spectra of trypsin-digested peptide fragments of neddylated BCR-ABL.**

The neddylation at K257, K500, K739, K802, K1025, K1135, K1590, and K1990 is respectively presented with the numbering at Lys residue. Detected peptide sequences are indicated in blue (b ions) and red (y ions)

Supplementary Figure 4

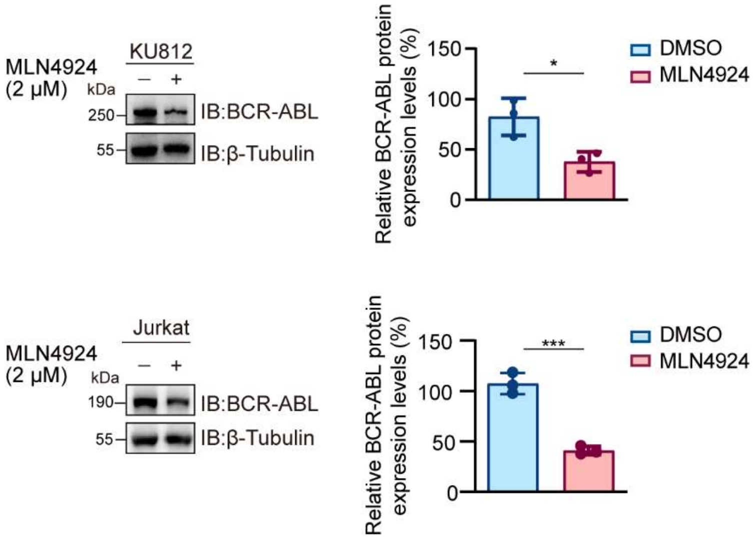


**Supplementary Figure S4. RAPSYN promotes BCR-ABL stabilization.** Immunoblotting analyses of BCR-ABL in KU812 and Jurkat cells treated with MLN4924 or DMSO for 24 h. Error bars, mean ± SD; * $p < 0.05$; *** $p < 0.001$; Student's t-test.

Supplementary Figure 5

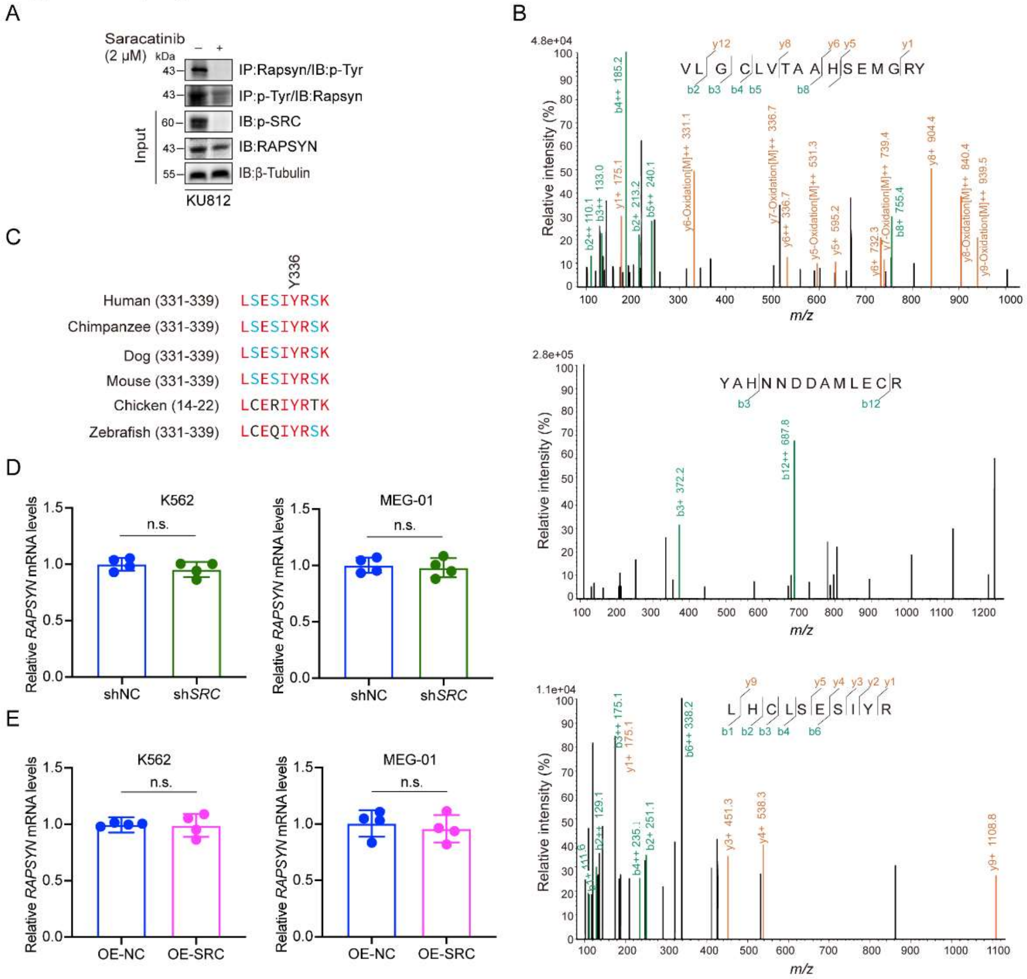


**Supplementary Figure S5. SRC-mediated phosphorylation at Y336 promotes RAPSYN stability.**

**A,** Assessment of RAPSYN phosphorylation in KU812 cells treated with saracatinib or DMSO for 24 h. **B,** LC-MS/MS spectra of trypsin-digested RAPSYN fragments (phosphorylated Y59, Y152, and Y336). The detected products are indicated by green (b ions) and orange (y ions). **C,** Sequence alignment of putative phosphorylated site Y336 from indicated species. **D,** Quantification of *RAPSYN* mRNA levels in K562 and MEG-01 cells transduced with sh*SRC* or shNC by RT-PCR. **E,** Quantification of *RAPSYN* mRNA levels in K562 and MEG-01 cells expressing exogenous *SRC* cDNA or corresponding empty vector by RT-PCR. *RAPSN* mRNA levels were normalized to that of *GAPDH* (D-E); error bars, mean ± SD; n.s., not significant; unpaired Student's t-test.

Supplementary Figure 6

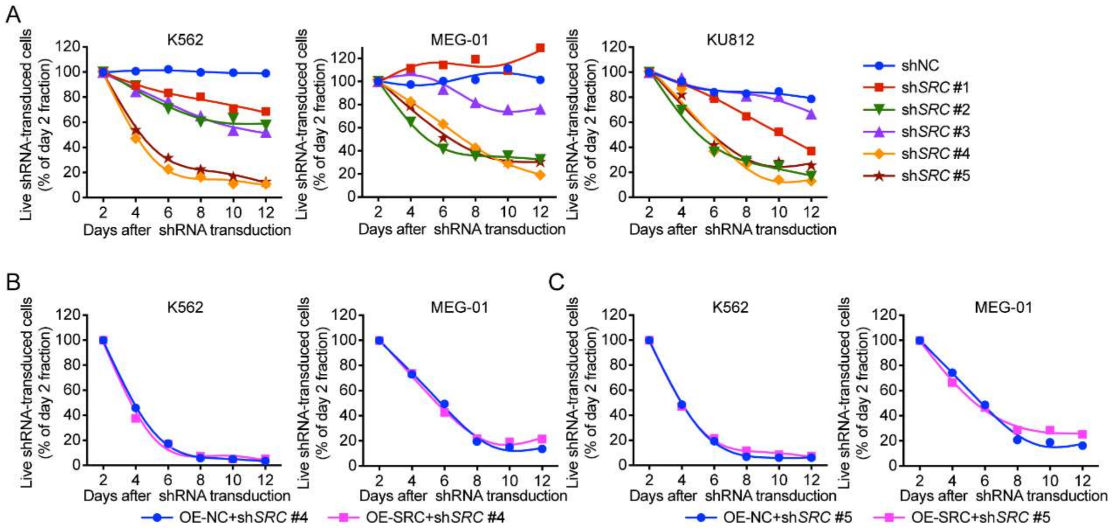


**Supplementary Figure 6. sh*SRC* #2 is a specific shRNA targeting the 3'UTR of *SRC*.**

**A,** Toxicity tests of all sh*SRC*s in Ph+ leukemia. **B-C,** Failed rescue of K562 and MEG-01 cells from sh*SRC* #4 (**B**) and #5 (**C**)-induced toxicity by exogenous expression of a *SRC* cDNA. Representative results from at least three independent experiments are shown (A-C).
